# Systematic evaluation of blastoid models of early human development

**DOI:** 10.1101/2024.07.11.603073

**Authors:** Hani Jieun Kim, Nazmus Salehin, Hao Huang, Xinran Zhang, Raja Jothi, Pengyi Yang

**Affiliations:** Computational Systems Biology Group, Children’s Medical Research Institute, Faculty of Medicine and Health, The University of Sydney, Westmead, NSW 2145, Australia; Embryology Unit, Children’s Medical Research Institute, Faculty of Medicine and Health, The University of Sydney, Westmead, NSW 2145, Australia; Epigenetics & Stem Cell Biology Laboratory, National Institute of Environmental Health Sciences, National Institutes of Health, Research Triangle Park, Durham, NC, 27709, USA; Charles Perkins Centre, School of Mathematics and Statistics, The University of Sydney, Sydney, NSW 2006, Australia; Kinghorn Cancer Centre and Cancer Research Theme, Garvan Institute of Medical Research, Darlinghurst, New South Wales, Australia

## Abstract

Recent generation of stem cell-derived blastoids that recapitulate the architecture and cellular constitution of the human blastocyst offers an experimental model for the elucidation of the biology of early human development. To evaluate the fidelity of the blastoids for modelling human blastocysts, we first established a reference map of cell identity and lineage differentiation of the human blastocysts through the integration and curation of single-cell transcriptome data of *ex vivo* blastocysts profiled at a range of developmental timepoints in culture. We next use this reference map to assess the coverage and the authenticity of cell lineages and the progression of lineage differentiation of various blastoid models generated with different cellular sources and protocols. This reference map generated from this study enables the benchmarking of blastoids that may guide the optimization of the protocol for the generation of high-fidelity blastoid models that faithfully recapitulate the natural human blastocyst.

## Introduction

The study of early human embryogenesis from the formation of the blastocyst and to gastrulation has been constrained by technical challenges and ethical concerns associated with human embryo research (Rossant and Tam, 2018; Rugg-Gunn et al., 2023). Recent achievements in generating blastocyst-like structures from stem cells, the blastoids, that are reminiscent of the human blastocysts in morphology and cellular composition has opened an avenue to glean knowledge of the biology of early human embryogenesis (Rossant and Tam, 2022). To date, a variety of protocols have been developed for generating human blastoids from various cellular sources such as naive pluripotent stem cells (PSCs) derived from reprogrammed embryonic stem cells (ESCs) (Kagawa et al., 2022, Yanagida et al., 2021, Yu et al., 2021), extended PSCs (EPSCs) (Fan et al., 2021), or iPSCs reprogrammed from fibroblasts (Liu et al., 2021). Nevertheless, varying levels of efficiencies of blastoid generation between protocols and limited extent of development of the blastoids in culture raise the reservation that these blastoids may not be high-fidelity models of the human blastocyst. Establishing a systematic evaluation framework for assessing the fidelity of the blastoid in modelling the human blastocyst is critical for enhancing the quality of this embryo model.

Here, we set out to establish a resource for a reference map of the human blastocyst and developed a computational workflow for assessing the quality of the blastoids generated by the current state-of-the-art protocols to recapitulate the attributes of the human blastocyst. To this end, we generated a comprehensive cellular identity reference map of the cell lineages and genetic programs of the developing human blastocyst by integrating single-cell transcriptomics profiling data of human embryos during the peri-implantation development of the *ex vivo* blastocyst (4 days post-fertilization, dpf; E4) to the equivalent of 12 dpf (E12) in culture (Molè et al., 2021; Petropoulos et al., 2016; Xiang et al., 2020; Zhou et al., 2019) (**Figure 1A**). This reference map sets the criteria for benchmarking the quality of blastoids generated in four protocols (Fan et al., 2021, Kagawa et al., 2022, Yanagida et al., 2021). Results of the benchmarking revealed the variable performance of blastoids in recapitulating the constitution and differentiation of cell lineages in the developing blastocyst. Collectively, our work provides a comprehensive and systematic evaluation of the developmental authenticity of *in vitro* cultured embryo models generated by the state-of-the-art protocols.

**Figure 1.**
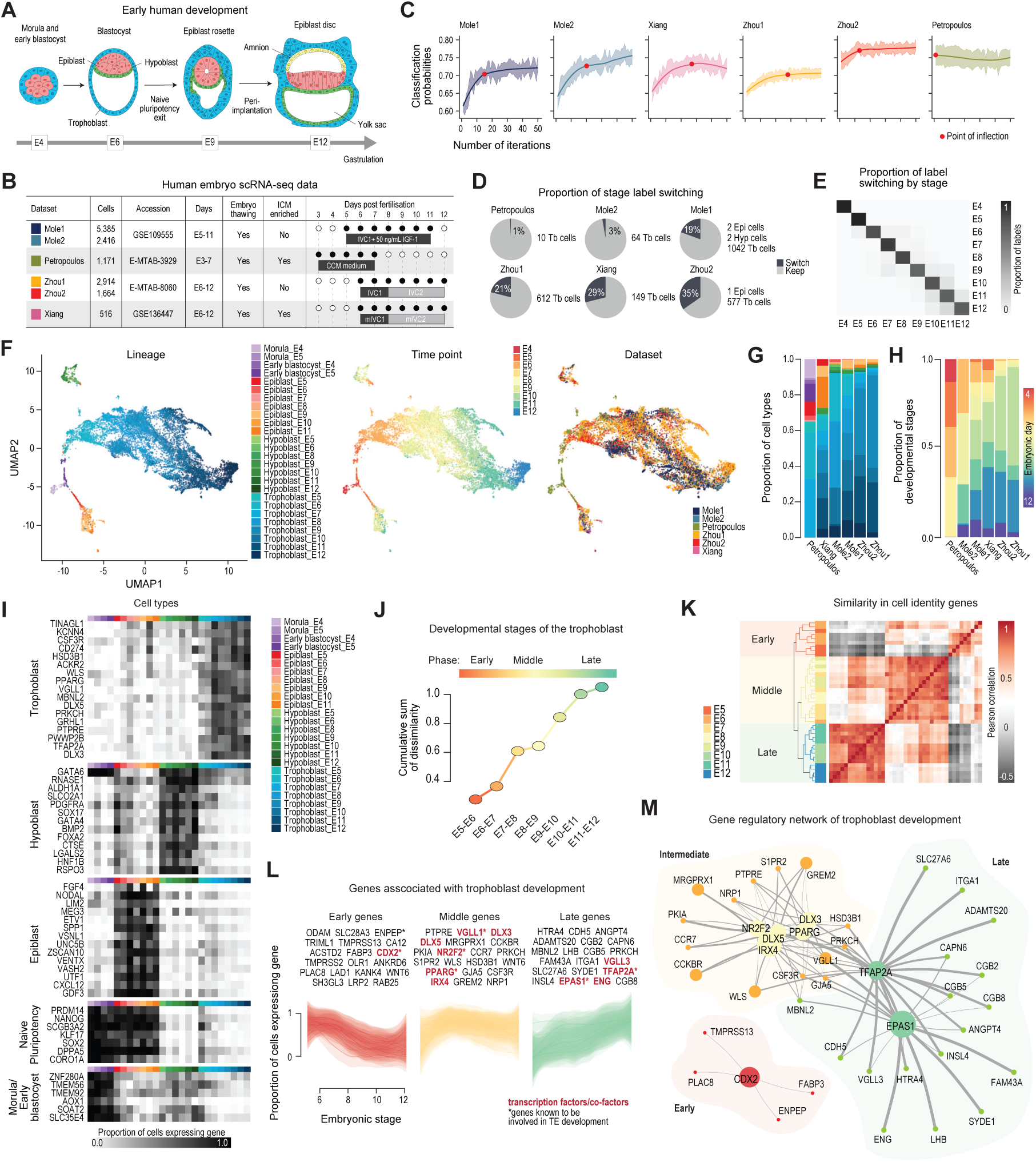
Generation of the in vivo embryo reference. **(A)** Schematic of early development of human embryo from embryonic (E) day 4 to day 12 post fertilization. **(B)** Single-cell RNA-seq datasets of ex vivo human blastocysts in culture for the collation of the reference map. Days: the span of equivalent stages of development (E: embryonic day) of ex vivo embryo in culture. TE, trophectoderm. **(C)** Scatter plot showing the improvement in classification probabilities (calculated using scReClassify) with each iteration in the iterative ensemble learning. The y-axis denotes the classification probabilities averaged across the classification probabilities from each cell type, where 1 denotes the highest accuracy. The final iteration (determined as the point of inflection) is calculated by fitting a generalised additive model across iteration and the probabilities. For each dataset, the point of inflection is indicated as a red dot. **(D)** Pie chart showing the proportion of cells of specific lineages in each dataset with switched developmental stage/time labels after the iterative ensemble learning. Tb, trophoblast; Epi, epiblast; Hyp, hypoblast. **(E)** A heatmap showing the proportion of cells switching by developmental stage/time. The rows denote time-label assignment of the cell using the original labels and the columns denote the new time-label assignment. The proportions are calculated from counts that are z-score standardised by each row. **(F)** UMAP representation of the integrated ex vivo embryo reference. The UMAPs are color-coded by lineage (left), time point (middle), and dataset (right). **(G)** Proportional bar plot of the synchronized cell lineage and developmental stage/time annotations for each dataset. The bars are color-coded by annotation labels (as in F). **(H)** Proportional bar plot of the developmental stage (timepoint) annotations for each dataset. The bars are color-coded by developmental stage (E4-E12). **(I)** Gene expression profiles of representative genes marking the morula/early blastocyst, naïve pluripotency, and the three major lineages: epiblast, hypoblast and trophoblast. The gene expression is calculated as the proportion of cells expressing the specific gene for each cell type (columns) averaged across the dataset. The grey-scale bar denotes the proportion: value 1 means all cells in the cell type are expressing the gene. Each column denotes a cell type documented in the reference (panels F, G). **(J)** Scatter plot showing the cumulative sum of dissimilarity (y-axis) between trophoblast cells at adjacent time points (x-axis). For the initial timepoint, the early blastocyst from E5 was used. The dissimilarity was calculated as the difference in correlation between two consecutive timepoints. The colors denote each pair of embryonic stages compared. **(K)** Pairwise correlation heatmap of the cell identity profiles of trophectoderm samples. The heatmap is clustered using hierarchical clustering and the terminal nodes of the tree is colored by embryonic timepoint. The heatmap is colored in terms of the degree in similarity calculated using the Pearson’s correlation where red and grey colors denote positive and negative correlation, respectively. **(L)** Smoothed scatter plot depicting the time-course expression profile of genes associated with trophoblast development, using aloess smoother to fit the curve. The x-axis denotes embryonic stage/time point and the y-axis denotes the expression level of genes quantified as the proportion of cells expressing the gene. The individual smoothed line denote the expression level of each gene listed above the plot. Transcription factors are highlighted in red and genes demonstrated to be involved in trophoblast development are marked by the asterisk. **(M)** Gene regulatory network of trophoblast development, compartmentalized by early (red), intermediate (yellow) and late stage (green) development

## Results

The first step towards benchmarking of embryo-like models is to build a comprehensive cell identity atlas of the human embryo. To establish such a reference, we developed a pipeline that iteratively learns from and integrates multiple human embryo single-cell RNA sequencing (scRNA-seq) datasets (Molè et al., 2021; Petropoulos et al., 2016; Xiang et al., 2020; Zhou et al., 2019) (**Figure 1B**). The timeline of development is inferred from the duration (days) in culture with the assumption that the chronology of *ex vivo* development is comparable to that *in vivo*. This pipeline consists of a two-step procedure that (1) cells are classified into one of five cell types (morula, early blastocyst, epiblast, hypoblast, or trophoblast) and then (2) their embryonic stage (time labels) are harmonizse across datasets (**Figure S1A** and see **Methods**). We first integrated the Petropoulos, Mole, and Zhou1 data that contain predefined labels using anchor-based integration through reciprocal PCA (Hao et al., 2021). We next performed annotation refinement using scReClassify, a semi-supervised learning method for assessing cell lineage label quality in the original annotation and if required corrects the cell label (Kim et al., 2019). This resulted in about 0.2% of cells being re-annotated into a different lineage, demonstrating the high quality of the original labels at the broad lineage level. Using this integrated data, we performed the label transfer onto Xiang and Zhou2 data, leading to a total of 14,250 annotated single cells of the *ex vivo* human embryos. We excluded cells from the pre-morula (83 cells) and the inner cell mass (14 cells) to retain the three lineages in the embryo from day 4 (E4) onwards.

**Supplementary Figure 1. Related to Figure 1.**
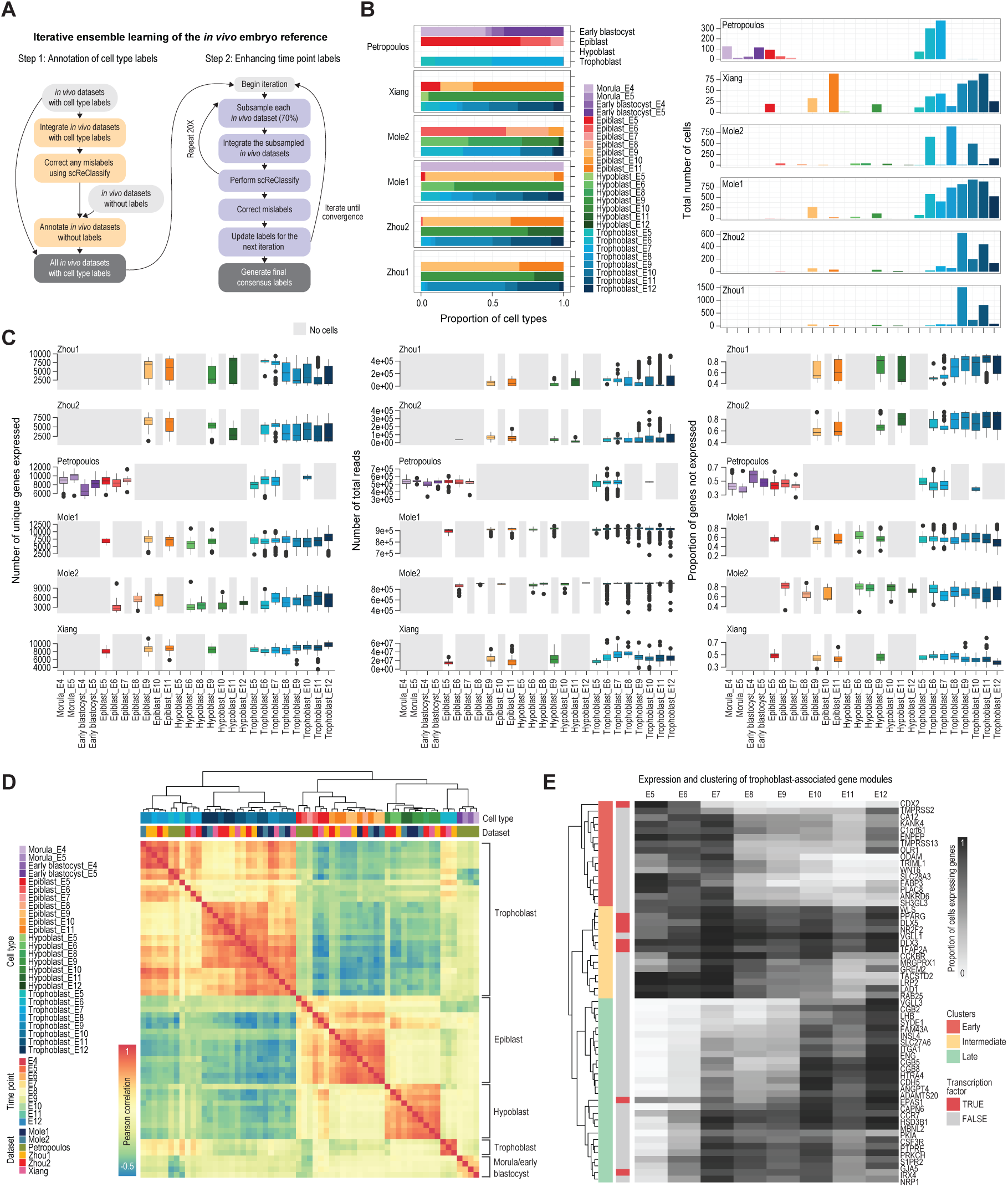
**(A)** Schematic of the iterative ensemble learning for optimising the cell type and time point annotations in the ex vivo embryo reference. **(B)** Proportion (left) and total cell count (right) of cell types found in each dataset. Bars are color-coded by the final cell type annotations. **(C)** Boxplot of three quality control metrics: number of unique gene expressed (left), number of total reads (middle), and proportion of genes not expression (zeros, right), for each dataset and cell type label. Boxes are color-coded by cell type as in panel B. **(D)** Pairwise correlation heatmap of cell identity scores calculated as the Cepo statistics for each dataset. The heatmap is clustered by the similarity in scores using hierarchical clustering. Cell types, time point and dataset are color-coded (left). Column: dataset by origin for each sample. **(E)** Gene expression profile of genes associated with the trophoblast development network. Gene expression is calculated as the proportion of trophoblast cells expressing the gene for each time point (columns) averaged across the dataset. The color bar denotes the proportion where value 1 means all cells in the cell type are expressing the gene. The rows of the expression heatmap are clustering using hierarchical clustering. The three clusters identified from hierarchical clustering are annotated by the trophoblast development (stage: early, intermediate, and late). Transcription factors are indicated in the color bar as red.

To correct for variability in the temporal progression of the *ex vivo* embryos cultured under different conditions (**Figure 1B**), we applied iterative ensemble learning to harmonise the temporal annotations between datasets (**Methods** and **Figure S1A**). scReClassify was applied to iteratively update the labels in the developmental stage/culture time point annotations in the integrated dataset until the annotations converged. Iterative synchronisation of the cell lineage annotations would improve model training to reach better classification probabilities, and an improvement in the cell type classification accuracy was observed with the iterative learning (**Figure 1C**). By fitting a generalised additive model and identifying the iteration at which the slope plateaus (**Methods**), a dataset-specific stopping point at which we deemed the cell type labels were sufficiently synchronised between datasets was determined. Most datasets (except the Petropoulos dataset) showed a 5-20% gain in classification probabilities, demonstrating a positive effect of the synchronisation of annotations (**Figure 1C**). Unlike other datasets, the Petropoulos annotations did not require any re-labelling, which might be explained by that these annotations were drawn from a recent re-annotation of these cell types (Meistermann et al., 2021). Approximately 20-35% of cells required synchronisation of the stage/time labels, with the majority of re-labelling for trophoblast cells (**Figure 1D**). As expected, most of the switching was between adjacent timepoints, with 76% of the re-labelling between E9-E12 embryonic stages, consistent with the anticipation that the asynchrony is entrained more at advanced stages (**Figure 1E**).

We visualised the results of the iterative ensemble learning by UMAP and confirmed visually the separation of cell types of the three major lineages across developmental timepoints (**Figure 1F**). Various quality metrics (such as the total number of cells per cell type, proportion of cell type, and per-cell metrics) were summarised for each dataset (**Figure S1B-C**). The Petropoulos and Xiang embryo datasets are depleted of the trophectoderm lineage, as this population may have been removed by immunosurgery prior to sampling (**Figure 1G**). In line with the timeline of embryo sampling and the duration of *in vitro* culture (**Figure 1C**), the majority of the earlier stage cells are found in the Petropoulos data, and cells of more advanced stages are found in the Xiang and Zhou data (**Figure 1H**). Importantly, we detected distinctive cell identity markers for the five major cell types, morula, early blastocysts, epiblast, hypoblast, and trophoblast (**Figure 1I**). Using the final cell lineage annotations, we generated the cell identity reference of the embryo using Cepo (Kim et al., 2021) for quantifying cell identity gene statistics for each cell lineage resolved by developmental timepoints, which are collated as a reference heatmap of the hierarchically clustered correlation matrix of the cell identity scores (**Figure S1D**). The results show that the three cell lineages broadly segregate into three groups, with the exception of the early-stage trophoblast that clusters with the early blastocyst and the morula, irrespective of dataset origin. Furthermore, we observed that the Cepo-derived cell identity scores enabled segregation of samples by developmental timepoints, as determined from the branching of the hierarchical tree (**Figure S1D**). These findings confirm the utility of Cepo to resolve genes that underlie the cellular identities in the embryo in a developmental stage/time-specific manner. The final cell identity reference of the embryo was derived by computing the average Cepo statistics by each lineage or lineage resolved by developmental stage.

To further corroborate our cell identity reference of the embryo, we exploited the large number of trophectoderm cells available in the reference map to tease out the developmental programs underpinning trophectoderm development (**Figure S1B**). By evaluating the dissimilarity between the cellular identity scores between adjacent developmental stages (see **Methods**), the trophectoderm- related samples can be grouped into three phases, with major changes in transcriptome occurring between E6 and E7 and between days E8 and E9 (**Figure 1J**). Consistent with previous findings (Lv et al., 2019), the categorisations are further confirmed by hierarchical clustering on the correlation matrix (**Figure 1K**), which shows that the three clusters equate to the separation of the early, intermediate, and late phases identified in **Figure 1J**. Visualisation of the gene modules of the three phases clearly revealed that the gene expression change follows the trend where genes in the early module that show a gradual decrease in the expression over developmental time comprise key markers of early trophectoderm development such as CDX2 (**Figure 2L and S2E**). The temporal profiles of genes in the intermediate module showed an increase in expression by E8 and persisted until the late stage, whilst genes in the late module showed a gradual increase in gene expression (**Figure 2L and S2E**). Of note, all late genes (except LHB and SLC27A6) and some of the intermediate genes (**Figure 2L**) were characterised as late genes in a previous study of 476 single-cell transcriptomes from trophectoderm across five developmental timepoints (E6-E10) (Lv et al., 2019). Finally, using these gene modules, we reconstructed the gene regulatory network that underlies the peri-implantation trophoblasts of the developing embryo (**Figure 1M**). Collectively, the analysis of cell identity genes and trophectoderm network analyses has generated a comprehensive integrated cell identity reference for benchmarking the experimental models of peri-implantation human embryo.

**Figure 2.**
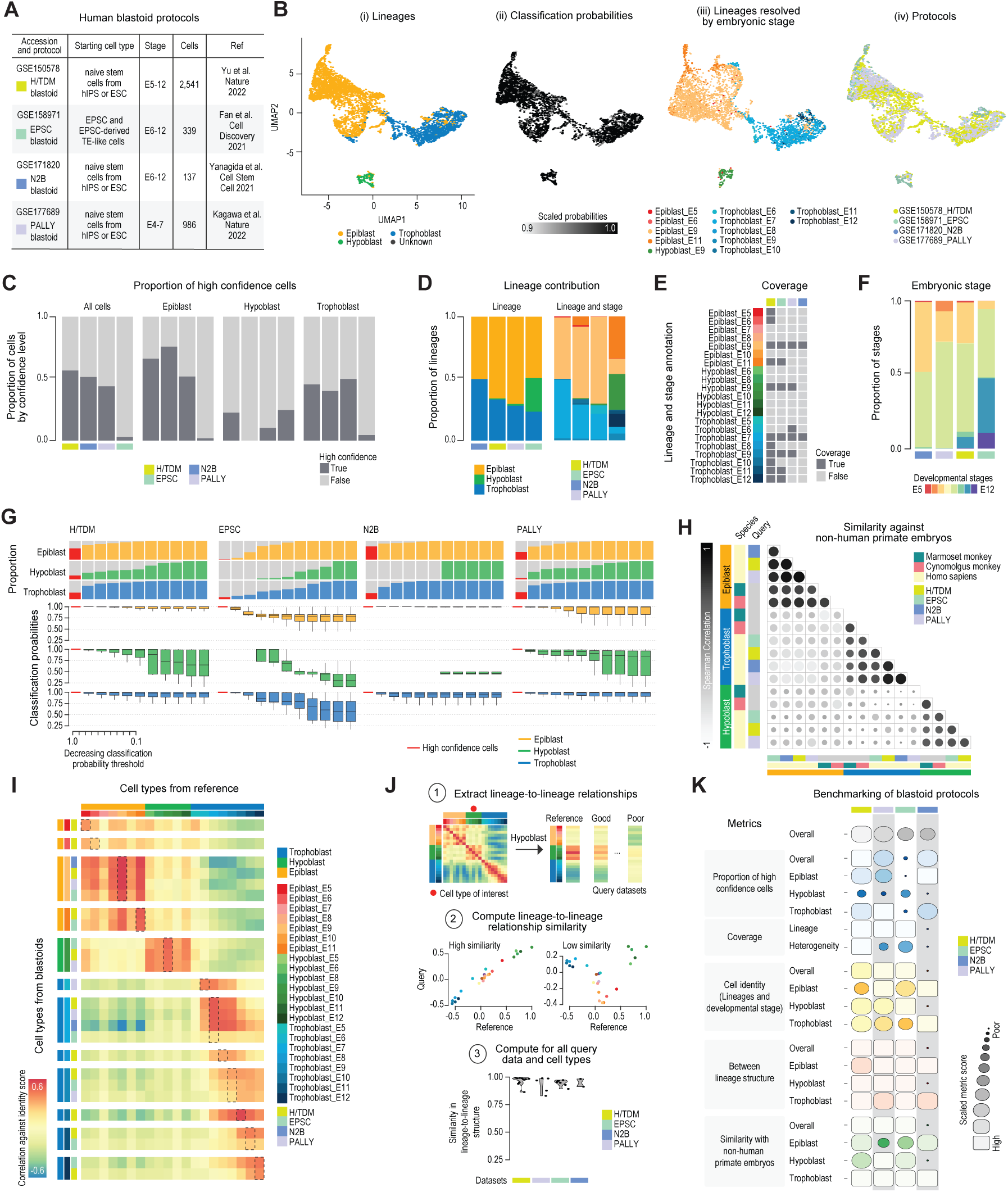
Benchmarking of in vitro blastoid protocols. **(A)** A summary of the single-cell RNA-seq datasets of the human blastoids. Stage: E5, E6 equivalent developmental timepoint at day 1 of culture, span of developmental stage inferred by days in culture. **(B)** tSNE representation of the integrated scRNA-seq datasets of the blastoids. The cells are color- coded by (i) cell lineage, (ii) classification probabilities scaled by each lineage group, (iii) cell lineage resolved by developmental stage and (iv) protocol. **(C)** Proportional bar plot of cells assigned as high (dark grey) or low (light grey) confidence for all cells and by individual cell lineage. The results are presented by dataset (horizontal axis, color-coded). **(D)** Proportional bar plot of the cell lineage (left) and lineage by developmental stage (right) annotations for each dataset color-coded as in (C). The bars colored by either lineage or lineage resolved by developmental stage. **(E)** Coverage of lineage types by developmental stage for the datasets. Present: dark grey), absent: light grey, column: data set, color-coded as in (C). **(F)** Proportional bar plot of the developmental stage annotations for each dataset, color-coded as in (C). Developmental stage: E4-E12, color-coded. **(G)** Boxplot plot showing the overall classification probabilities (y-axis) for cell groups positioned against a deceasing classification probability threshold (x-axis). The red bar denotes the cell group containing the high confidence cells used in the benchmarking analysis. Plots are color-coded by cell lineage. The results are visualised for each dataset separately. The proportional bar plots denote the proportion of cells from cell group (separated and color-coded by lineage as in the boxplots) contributing to the total number of cells in the lineage. **(H)** Pairwise Spearman correlation indices of cell identity scores of cell lineages in blastoids generated by different protocols (color-coded) and in non-human primate embryos (marmoset [turquoise] or cynomolgus [pink] monkeys). The heatmap is clustered using hierarchical clustering on the correlation matrix. **(I)** Heatmap of Pearson’s correlation of cell identity scores between the reference (columns) and the cell lineages of the blastoids (rows). The heatmap is ordered and color-coded by lineage (epiblast, hypoblast, and trophoblast) and developmental stage/time in culture. Within each lineage and developmental stage, the rows are ordered by the strength of correlation, where the datasets (color-coded) are ordered from highest to lowest correlation. **(J)** Steps of analytics workflow. 1: Identity lineage-to-lineage relationships from the reference to evaluate fidelity of this relationship in the query blastoid datasets that may show varying capacities to recapitulate the reference. 2: Taking a hypoblast sample as an example, the scatterplots exemplify a query dataset that shows a high concordance in the lineage-to-lineage relationships (left) and a low concordance relationship (right). 3: A violin plot showing overall distribution of the lineage-to-lineage structure similarity performance for each dataset, color-coded as in (I). **(K)** The benchmarking results of the blastoids (column by protocol) on five criteria: proportion of high confidence cells, coverage, cell identity, lineage-to-lineage structure, and similarity with non-human primate embryos. The scaled metric scores are depicted as squircles where the size denotes the score from large=high, to small=low for each row. Squircle sizes are comparable within each metric class but not between metric classes. Blastoids that performed well have larger squicles.

The recent generation of the stem cell-based blastocyst-like structures, the blastoids, breaks new ground for modelling early human development (Fan et al., 2021; Kagawa et al., 2022; Liu et al., 2021; Xiang et al., 2020; Yu et al., 2021). To evaluate the quality of these blastoids generated these protocols, we collated public datasets obtained from some of the protocols which varied in the starting cell type and the specific media compositions, addition of supplements or growth factors, and the duration of culture of the initial culture, aggregation and blastoid differentiation conditions (**Figure 2A**, **Table S1**). We developed an evaluation framework by using our cell identity reference of the human embryo (**Figure S1D**) and a panel of metrics for distinguishing the level of fidelity of different blastoids in recapitulating the *ex vivo* human embryos. We implemented five major classes of metrics: 1) proportion of high confidence cells, 2) cell type coverage, 3) cell identity of each lineage, 4) lineage-to- lineage structure, and 5) similarity with the non-human primate embryo. The full description of how the references were generated and the metrics are computed can be found in the **Methods**.

**Supplementary Table 1.**
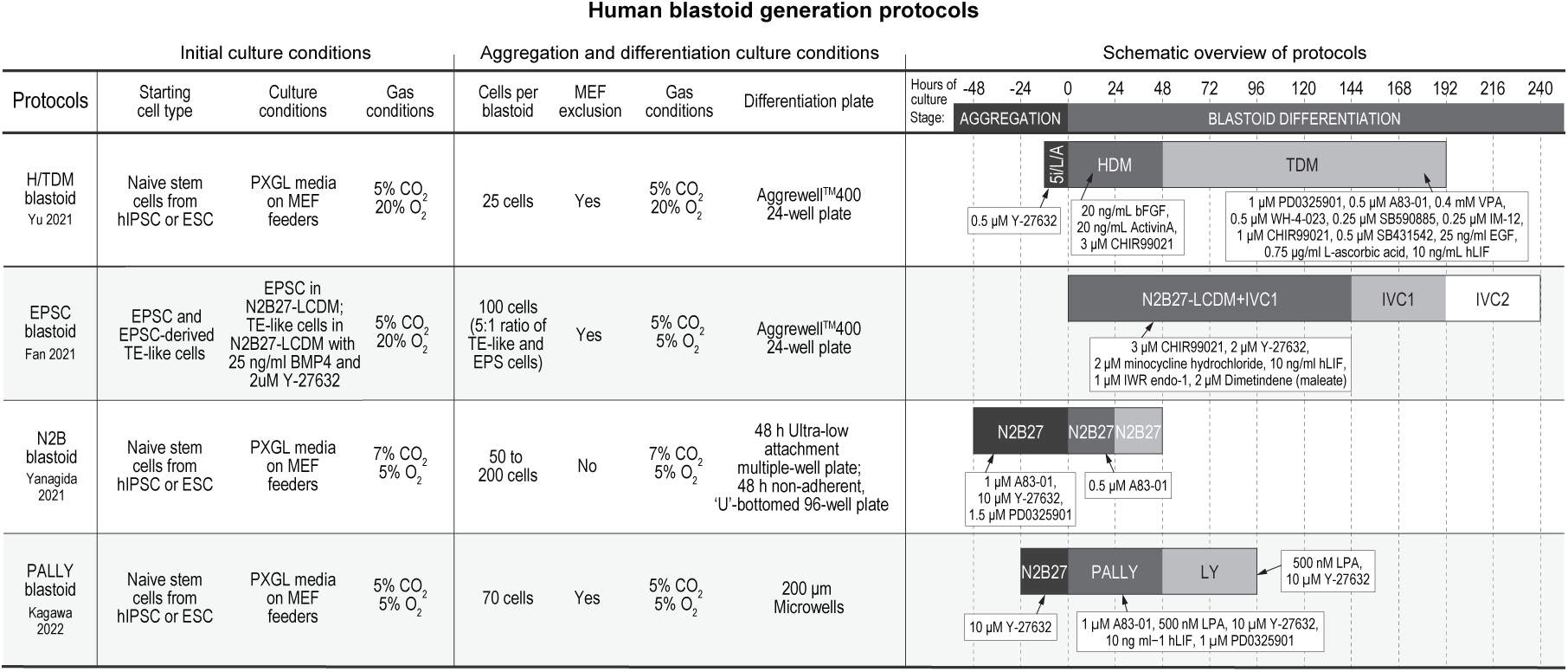
A summary table of human blastoid generation protocols. Abbreviations: human induced pluripotent stem cells (hiPSC), embryonic stem cells (ESC), expanded pluripotent stem cells (EPSC), trophectoderm (TE), mouse embryonic fibroblasts (MEF), hypoblast differentiation medium (HDM), trophoblast differentiation medium (TDM), *in vitro* culture (IVC).

To accurately identify the cell states of the blastoids, we classified single cells from blastoids into one of three lineages of the early human embryo (epiblast, hypoblast, or the trophoblast) using our embryo reference and scClassify, a hierarchical cell type classification method (Lin et al., 2020). Any cells that were not classified to any of the three lineages were labelled as unknown. Using the classification probabilities, we identified cells that have been assigned a cell type with high confidence (**Figure S2A-C**, see **Methods**). Next, a finer resolution scClassify model was then built to predict the developmental stage of the assigned cells. In particular, we built a model for each lineage using the lineage and developmental stage resolved cell type annotations. Any cell labelled as an unknown or low confidence assignment was excluded from the classification of developmental stage. For each classification result, the cell type labels were determined by the majority voting from individual classification labels using each of the *ex vivo* reference datasets. The final classification results were visualised in **Figure 2B**.

Our multiscale classification results revealed a large portion of low confidence cells with most protocols (H/TDM, N2B, and PALLY) showed approximately 40-55% of high confidence cells and a substantial portion of low confidence cells (**Figure 2C**). Breaking down by proportion per lineage, we observed that H/TDM, N2B, and PALLY protocols contain the highest proportion of epiblast-like cells, followed by trophectoderm-like and the hypoblast-like cells (**Figure 2C**). Whilst these protocols generated single cells with similar quality control metrics (**Figure S2E**), the blastoids generated from the EPSC protocol demonstrated the lowest proportion (5%) of successfully allocated cells according to our embryo atlas, which may be partially explained by the significantly higher number of cells sampled in this study (**Figure S2D**). N2B blastoids contained 50% unknown cells, which may be off-target cells that depart from the three lineages. Collectively, these findings show that most blastoids have on average 50% of the cells being highly confident embryonic cells, with H/TDM and PALLY blastoids showing a higher proportion of confident cell types.

We defined cell type coverage in terms of two metrics: the capacity of contributing to the three lineages and of recapitulating the heterogeneity of cell states. We measured the capacity to generate the three lineages, each with five or more highly confident cells. We found that most blastoids were capable of generating all three lineages except the N2B blastoids (**Figure 2D**). As a metric to assess heterogeneity, we further counted the number of developmental stages covered by the lineage. In view of the extended period of culture (between 48 to 240 hours), we made an assumption that the blastoids will display some degree of asynchronous development, leading to the capture of heterogeneous cell states associated with the progression of lineage differentiation. As anticipated, we observed that the blastoids with more extended blastoid differentiation culture systems (H/TDM and EPSC blastoids) generated more heterogeneous cell states and a greater proportion of cells associated with an advanced developmental stage than blastoids (N2B and PALLY blastoids) developed for a shorter duration in culture, which contained a greater proportion of cells associated with the early phase of lineage differentiation (**Figure 2E-F**).

We next defined two metrics for the measurement of cellular identities: 1) a metric that quantitatively measures the similarity of a lineage of interest to the corresponding lineage in the reference and 2) a metric that measures the recapitulation of relationships between a lineage and other lineages (**Methods**). These two metrics provide two views of cellular identity in the transcriptome data. For the lineage-specific cell identity metric, we performed the analysis at two layers of annotation (lineage only and lineage at developmental stages). At the coarser level of annotation, we found that the H/TDM and PALLY blastoids generated cells with the highest probabilities across the three lineages (**Figure 2G**). Whilst the N2B blastoids generated cells with comparable probabilities to the H/TDM and PALLY protocols, this protocol failed to generate any high confidence hypoblast-like cells. Whilst the EPSC blastoids did generate epiblast-like cells that were highly confident, the proportion of cells that passed the threshold was low (**Figure 2C, 2G, and S2D**). At the finer level of annotation, we evaluated the results in terms of their correlation to the cell lineages in the reference. The findings mirrored the results observed in Figure 2G and are broken down into the developmental stages for each cell lineage (**Figure 2I and S2J**).

**Supplementary Figure 2. Related to Figure 2.**
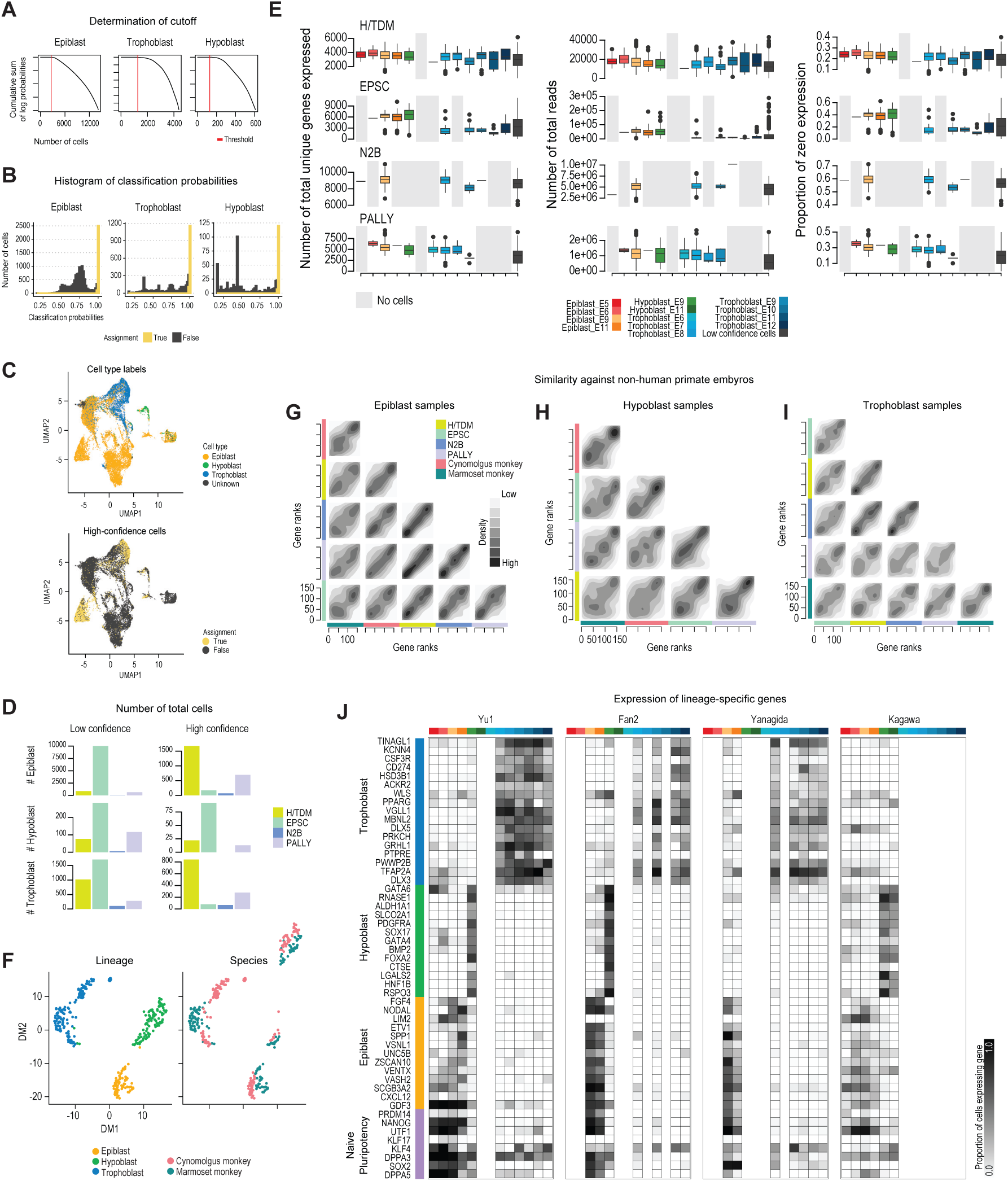
**(A)** Plot of the cumulative sum of the log probabilities of classification for each cell lineage. Cells are ordered by decreasing classification probabilities across the x-axis. Y-axis denotes the cumulative sum of the log probabilities of classification. The red line denotes the point of inflection which is defined as the first instance at which the difference between two elements in the sequence is greater than 0.01. **(B)** Histogram of the classification probabilities for each lineage. Classification probabilities are min-max scaled by lineage. Histograms are colored by whether the cells have passed the threshold defined in (A). Cells that have passed the threshold (received a TRUE assignment and are deemed highly confident) are plotted in the yellow histograms, and those that failed (received a FALSE assignment) are plotted in the grey histograms. **(C)** tSNE representation of the blastoid scRNA-seq datasets. Cells are colored by assignment of confidence where yellow denote high confidence cells and grey denotes low confidence cells. **(D)** Bar plot of total cell numbers of low (left) and high (right) confidence cells by lineage (rows) and dataset (color-coded). **(E)** Boxplot of three quality control metrics, number of unique gene expressed (left), number of total reads (middle), and proportion of zeros (right), for each dataset and cell type label. Boxes are color-coded by cell type. **(F)** tSNE representation of the integrated non-human primate embryo reference. The cells are color-coded by cell lineage (left) and species (right). **(G-I)** Pairwise contour plot of ranked cell identity scores calculated as Cepo statistics for each dataset (color-coded) and lineage: (G) epiblast, (H) hypoblast, and (I) trophoblast. Higher density of genes is denoted by a darker contour. **(J)** Expression profiles of representative genes marking cells with naïve pluripotency, and the three major lineages (epiblast, hypoblast, and trophoblast). The gene expression is calculated as the proportion of cells expressing the genes for each cell lineage across the developmental stage (columns) for each dataset. The grey-scale bar denotes the proportion of cells expressing the gene where a value 1 means all cells in the cell lineage are expressing the gene. Each column in a heatmap denotes a cell lineage by stage, color-coded as in (E).

Unlike the lineage-specific cell identity metric that quantifies how well a given cell type found in the embryo reference is recapitulated in the blastoids, the second metric provides information on how well the relationships between a cell type with another cell type is captured. We called this metric “lineage-to-lineage similarity” to dissect how well the between-lineage structure in the embryo reference is imitated by that in blastoids (**Figure 2J**), thus providing a measure of similarity on a cross-lineage scale that is distinct to the lineage-specific measures of the first metric (see **Methods**). We found that whilst most protocols performed well in terms of this metric, the H/TDM blastoid performed particularly well. We believe this metric will be a useful metric that can discriminate blastoid cells that despite their assignment to the reference with high confidence may demonstrate a low recapitulation of the lineage-to-lineage structure, identifying the presence of aberrant between lineage transcriptomics profiles that are independent from the lineage-specific ones.

Because of their close phylogenetic relationships and similarities to humans, non-human primates (NHP) are amenable models for embryo research where there are barriers to access human embryonic material. As the final metric, we evaluated whether the human blastoids display resemblance of lineage profile to that of NHP embryos. We integrated transcriptomics data from two NHP primates, the Cynomolgus and the Marmoset monkeys (Boroviak et al., 2018; Nakamura et al., 2016), leading to a collection of 72 epiblast, 107 hypoblast, and 147 trophectoderm cells (**Figure S2F**). Using the same lineage-specific cell identity metric with genes prioritised on the basis of the cell identity genes of the NHP embryos (see **Methods**), we computed the similarity between the cell identity statistics of the human blastoids and those of the NHP embyors (**Figure 2H and S2G-I**). We found that the samples were clearly segregated in terms of their lineage of origin irrespective of species (**Figure 2H**). These findings demonstrate that many lineage-specific genes are conserved across monkeys and humans and further support the usefulness of the human embryo reference to identify the three cell lineages in blastoids.

Finally, we summarised the results of these benchmarking metrics for the four human blastoids (**Figure 2K**). The performance was variable across the blastoid types with certain blastoids performing well for one lineage over the others. The better performers were the H/TDM and PALLY blastoids. Both were able to generate the three lineages that robustly recapitulate the cellular identities scores of the *ex vivo* embryo. The N2B blastoids demonstrated a specific depletion of hypoblast-like cells, where none of the cells classified as hypoblasts were of high confidence. The epiblast-like and trophoblast-like cells generated in the N2B blastoids showed comparable scores to the top performance blastoids across the metrics. Lastly, we found that the EPSC blastoids generated the least proportion of high confidence cells, yet the high-confidence cells matched well with the *ex vivo* embryo reference. In conclusion, this characterisation of the blastoids provides a measure of the fidelity of the blastoid in modelling the human blastocyst. We anticipate that the *ex vivo* embryo reference would be a useful data resource for the benchmarking of future models of blastoids and other integrated embryo models.

## Discussion

Whilst a key innovation of this study is the generation of the *ex vivo* embryo reference and the benchmarking of human blastoids generated by differentiation protocols against this embryo reference, one crucial limitation is that the metrics alone cannot be used as a measure of the developmental competence whether that be implantation or extended post-implantation development because none of the blastoids have been tested for these developmental outcomes (Heidari Khoei et al., 2023). Attempts at the prolonged culture of human blastoids have so far been few, with studies demonstrating that blastoids fail to develop for more than a few days when attached to tissue culture plastic or to monolayers of endometrial cells, an environment that mimics implantation *in vivo* (Kagawa et al., 2022; Yanagida et al., 2021; Yu et al., 2022). Clearly, improved culture systems that better mimic the implantation niche are requisite for maximising the research opportunities opened up by integrated embryo models. Another limitation of this study is that the fidelity metrics do not address important features of the embryo such as the profiles of the other omics layers, spatial patterning, and domain-specific characteristics. Recently, a few studies have begun profiling the primate embryo using spatial single-cell technologies (Bergmann et al., 2022). Such multi-omics studies, together with experimental studies of developmental competence, will expand not only the repertoire of metrics that can be used to interrogate human embryogenesis but also to pinpoint the missing casual links that make certain blastoids perform better than others.

## Acknowledgments

We thank our colleagues at the Children’s Medical Research Institute for their constructive feedback. This work is supported by a National Health and Medical Research Council (NHMRC) Investigator Grant (1173469) and a Metcalf Prize to P.Y.

## Author contribution

H.J.K. and P.Y. conceived the project; H.J.K. generated the references and developed the benchmarking framework with support from P.Y. and X.Z.; H.J.K. analyzed the data with support from P.Y., N.S., and H.H.; and H.J.K and P.Y .wrote the manuscript. All authors reviewed, edited, and approved the final version of the manuscript.

## METHODS

### QUANTIFICATION AND STATISTICAL ANALYSIS

#### Public single-cell RNA-seq datasets of the human embryos and blastoids

##### Data collection and initial processing

The raw gene by cell count matrices of scRNA-seq datasets were retrieved from the NCBI Gene Expression Omnibus (GEO) unless otherwise stated. All datasets are related to the human pre- and peri-implantation blastocyst or *in vitro* blastoids. The raw count or RPKM matrices downloaded from the public portals were library-size normalised and log-transformed using the *logNormCounts* function of the scater R package (McCarthy et al., 2017) or the *NormalizeData* function of the Seurat R package (Hao et al., 2021) using the default settings. The cell lineage and developmental stage annotations from the original studies were used when available.

##### Data filtering

We filtered out suboptimal cells in the reference data by using the *isOutlier* function from scran (Lun et al., 2016), which identifies outliers by setting a threshold based on the metric of deviation from the median. In particular, cells are filtered on the basis of (i) the number of expressed genes with a threshold of 2 deviation away from median and (ii) the total number of counts in the sample with a threshold of 3 deviation away from median. A total of 300 cells carrying too few or too many counts and genes were removed. We further removed potential doublets in each biological batch by using the DoubletFinder R package following their default settings (McGinnis et al., 2019). Specifically, the doublet detection rate was set to 0.05 leading to the identification of 773 potential doublet events. Collectively, a total of 1,024 out of 15,444 samples considered as either doublets or poor quality cells were filtered from the data prior to all downstream analyses. Finally, for each dataset, genes that were not expressed in any of the cells were removed.

#### Generation of the integrated *in vivo* human embryo reference

##### Iterative ensemble learning to optimise cell lineage annotations of the in vivo embryo reference

To establish a high-quality cell identity reference of the human embryo, we devised a framework to harmonise the cell lineage and developmental stage annotation of datasets generated by different studies so as to maximise the information gain from the resulting *in vivo* reference (**Figure S1A**).

For the cell lineage annotation, we performed the classification of unlabeled cells by generating an initial reference from the three datasets (Petropoulos, Mole, and Zhou1) with pre-annotated cell lineage labels (Molè et al., 2021; Petropoulos et al., 2016; Zhou et al., 2019) using an anchor-based integration approach (Hao et al., 2021). Briefly, This procedure requires three broad steps: dimension reduction using reciprocal PCA or canonical correlation analysis, identification of anchors between pairs of datasets using these reduced dimensions, and integrating datasets using the integration anchors. The pairwise correspondences between individual cells from each dataset are considered to be from the same biological state and can be used to harmonise and transfer information between the two sets. We integrated the three datasets with pre-annotated cell lineage labels using reciprocal PCA with 50 dimensions. For the selection of anchors, a total of 5 neighbours and 2000 features were used. Using the integrated and normalised expression values, we performed PCA analysis to generate 50 principal components. Using these components, we then performed post-hoc re-labelling with scReClassify (Kim et al., 2019). scReClassify is a semi-supervised learning method for assessing cell lineage label quality in the original annotation from the scRNA-seq datasets. We applied the random forest classifier with 10 ensemble learners. Only cells that received probabilities of an alternative assignment of greater than 0.8 (i.e., when more than 8 out of 10 learners identify a cell-type label as mis-labelled) were re-annotated. Finally, using the integrated initial reference with optimised cell-type annotations, we performed label transfer to the other unannotated *in vivo* datasets (Xiang et al., 2020; Zhou et al., 2019) using *FindTransferAnchors* and *TransferData* functions from the Seurat package. PCA was projected from the initial reference onto an unannotated dataset for which 50 and 30 PCs were computed on the two sets, respectively. The cell pairwise anchor correspondences between the initial reference and query were used to transfer the lineage annotations from the reference to the query cells. Importantly, we ensured that the cell lineage label transfer retained the developmental stage annotations of the reference cells, allowing the query cells to inherit both the lineage and developmental stage labels. Together, this procedure leads to a reference data that integrates five individual datasets.

We reasoned that further synchronisation of the developmental stage annotation is required given that the blastocysts are cultured *ex vivo* for varying periods of time by different laboratories using different culture protocols prior to capturing their transcriptomes through scRNA-seq. Such technical variation may lead to variability in the progression of lineage development and hence inconsistent developmental stage annotations between datasets, which may be attributable to annotation of cells purely on the basis of the number of days cultured. Therefore, we developed an approach that synchronises the developmental stage annotation (which might arise from asynchronous differentiation of cells between different cell culture conditions) through an iterative ensemble learning procedure using scReClassify. In particular, an iteration consisted of the following steps: (i) randomly subsampled the *in vivo* datasets to 70% of the total cell number; (ii) integrated the subsampled references using an anchor-based integration and canonical correlation analysis approach (Hao et al., 2021); (iii) using the integrated and normalised expression values, performed PCA analysis to generate 50 principal components; and finally (iv) performed synchronisation of the developmental stage annotation using the scReClassify framework as described above. This procedure was repeated 20 times. To ensure an accurate synchronisation, we corrected a developmental stage label of a cell if the re-annotation of the label occurred in more than 80% of the training (i.e., at least 16 out of 20 times), thus taking a conservative approach to update the developmental stage labels. At the start of each iteration, the annotations were updated with those from the previous iteration. We performed this procedure iteratively until the scReClassify probabilities plateaued at which point we deemed there was limited gain in the ensemble learning and that the cell type labels were sufficiently synchronised between datasets.

##### Identification of the point of inflection of the scReClassify probability curves

We developed an approach to dynamically identify the optimal number of iterations required for each dataset for its harmonisation and developmental stage synchronisation. Because some datasets may require only minimal improvement in annotations, whilst others may require several iterations before the benefits of the ensemble learning plateaus, the optimal number of iterations was determined as the point of inflection of when the scReClassify probabilities begin to plateau, indicating a slowing down of gain in benefit from the ensemble learning. We fitted the generalised additive model (GAM) (Hastie, 1992) using the iterations as the predictor and the scReClassify probabilities as the response variable, where the latter was assumed to follow a normal distribution. We calculated the scReClassify probabilities for the major lineages separately and used the average probabilities in the model fitting. The thin plate regression spline smooth function was applied. Then we evaluated the slope by calculating the first derivative (i.e., the slope) of the GAM spline. We defined the plateau (i.e., the point of inflection) as the first instance in which the slope is not significantly different from zero. The iteration equating to the inflection point, rounded to the nearest integer, was used to extract the final labels for each dataset.

#### Identification of lineage-specific developmental stage-resolved markers of the early blastocyst

##### Derivation of cell identity gene statistics and the embryo cell identity reference

To derive the cell identity gene statistics for the human embryo atlas, the normalised and log-transformed cell by gene count matrix were analysed using the Cepo package (Kim et al., 2021) for quantifying cell identity gene statistics for each cell lineage resolved by developmental stage based on the differential stability metric. The final cell identity reference of the embryo was derived by computing the average Cepo statistics by each lineage or lineage resolved by embryonic stage.

##### Construction of the gene regulatory network depicting trophectoderm development

To construct a gene regulatory network that underlies trophectoderm development, we first categorised the trophectoderm samples into distinct phases in terms of their development. To unbiasedly identify the distinct phases of development, we calculated the cumulative sum of dissimilarity using the cell-type-specific identity scores derived using Cepo. By comparing the change in dissimilarity between two adjacent timepoints, we determined three distinct phases whereby the change in sum dissimilarity is smaller than 0.1. Hierarchical clustering on the cell identity profiles further confirmed the grouping of the trophectoderm samples into the three distinct phases, which we defined as “early”, “intermediate”, and “late”.

Next, we extracted the genes identified as associated with trophectoderm identity and development by taking the top 20 genes on the basis of the Cepo statistics for each trophectoderm sample from the embryo reference. Using these genes, we constructed a gene regulatory network using the igraph R package with a layout calculated by using Fruchterman-Reingold layout algorithm (Csardi and Nepusz, 2005). The temporal mRNA expression of a gene *x* was first standardised across time points using z-score transformation:

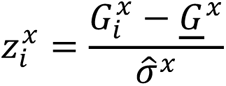

where for each gene 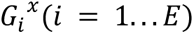 is the expression level of its mRNA at an embryonic timepoint *i*, *E* is the number of developmental stages included in the data, and *G^x^* and *σ^x^* are the average expression and variance across the timepoints. Using the standardised values, we assigned target genes for each transcription factor identified by calculating the strength of association between transcription factor expression and target gene expression. We assigned a transcription factor to a target gene if the association (computed using the Spearman correlation) was greater than 0.8. Any associations below 0.8 were discarded. The final gene regulatory network was weighted at two levels: (i) the nodes were weighted in terms of the total number of edges connected to the node, where a larger node denotes a greater number of links, and (ii) the edges were weighted in terms of the strength of the association, where a thicker connection denotes a stronger transcription factor and target gene link.

#### Protocols for generating human blastoids

##### Brief description of the blastoid protocols

1. **GSE150578_H/TDM protocol** (Yu et al., 2021). Naive pluripotent stem cells (hPSCs) are grown on mouse embryonic fibroblast feeders in 5i/L/A or PXGL medium. To form a blastoid, around 25 hPSCs are placed in an AggreWell^TM^400 24-well plate and incubated for 12 hours until aggregates form. The cells then undergo a two-stage differentiation process, switching from the PSC culture medium to hypoblast differentiation medium (HDM) and then to trophoblast differentiation medium (TDM) after 48 hours, which is refreshed every two days. After six to eight days of TDM culture, human blastoids can be observed.
2. **GSE158971_EPSC protocol** (Fan et al., 2021). The study generates blastocyst-like structures, called EPS-blastoids, using extended pluripotent stem (EPS) cells. The protocol uses a 3D, two-step differentiation protocol. First, EPS cells are converted from induced pluripotent stem cells using a previously described protocol (Yang et al., 2017) and cultured in human N2B27-LCDM medium on MEF feeders. To induce blastoids, the human EPS cells were differentiated in N2B27-LCDM medium supplemented with BMP4 for three to four days. Then, the BMP4-treated trophectoderm-like cells are mixed with EPS cells at a 5:1 ratio and seeded onto a AggreWell^TM^400 24-well plate pretreated in the human EPS-blastoid medium (EPS and IVC1 medium mixed at a ratio of 1.5:1 [v/v]). The culture medium was replaced every day. After 6 days of culture, blastoids are collected for further analysis, including *in vitro* culture.
3. **GSE171820_N2B protocol** (Yanagida et al., 2021). Naive pluripotent stem cells (hPSCs) are grown on mouse embryonic fibroblast feeders in PXGL medium. To form blastoids, 50-200 hPSCs are first seeded onto a ultra-low attachment multiple-well plate in PD+A83+Y medium. After 42-48 h, cell aggregates are manually transferred using mouth pipetting to an non- adherent, ‘U’-bottomed 96-well containing N2B27 medium supplemented with A83-01 to induce blastoid differentiation. At day 3 of culture, the blastoid is manually transferred into N2B27 medium without A83-01 and then collected at day 4 for further analysis.
4. **GSE177689_PALLY protocol** (Kagawa et al., 2022). Naive pluripotent stem cells (hPSCs) are grown on mouse embryonic fibroblast feeders in 5i/L/A or PXGL medium. To form a blastoid, approximately 70 hPSCs are placed in 200 µm diameter 96-well Microwell plates in N2B27 medium containing Y-27632 and incubated for 24 hours until aggregates form. Then, the cells then undergo a two-stage differentiation process: The cells are cultured in PALLY medium for two days (with the medium refreshed every 24 h) at the first stage. The medium is replaced with N2B27 medium supplemented with LPA and Y-27632 for another two days, completing the blastoid differentiation at 96 h of culture.

A schematic summary of the blastoid protocols evaluated in this study is also provided in **Table S1**.

#### Integration and classification of human blastoid scRNA-seq datasets

##### Embedding blastoid transcriptomes into a shared latent space

To embed the single-cell transcriptomes into a shared latent space, for each batch the count matrix was first normalised to the total number of reads and then multiplied by a 10,000 scaling factor. Then the top 2,000 highly variable genes determined through variance stabilising transformation were prioritised by their variance across all the scRNA-seq batches. Next, the cell pairwise anchor correspondences between different single-cell transcriptome batches were identified with 30- dimensional embeddings from reciprocal principal component analysis (Hao et al., 2021). Using these anchors, the scRNA-seq datasets were integrated and transformed into a shared space. Gene expression values were scaled for each gene across all integrated cells and used for principal component analysis (PCA). For the identification of anchors and integration of the blastoid datasets, *k. anchor* and *k.weight* were set to 5 and 50, respectively. To generate the two-dimensional visualisations of the single-cell transcriptomes derived from blastoids, the single cells were projected into tSNE space by using the first 50 principal components (PCs) generated on the integrated normalised matrix.

##### Multiscale classification of blastoids

To accurately identify the corresponding cell states of the early human embryo in the blastoids, we performed multiscale classification that consists of three broad stages: (i) classification of the three major lineages; (ii) identification and filtering of high confidence cells; (iii) and finally the classification of cells to their developmental stages with respect to the human embryos.

We performed cell type classification using the scClassify framework (Lin et al., 2020) that incorporates multiscale learning. First, we built a model from references with cell type annotations of the three main lineages of the human embryo (i.e., epiblast, hypoblast, or the trophoblast). The cell lineage labels were determined by the majority vote from individual classification labels using each of the *in vivo* reference dataset. To enable the majority of cells to be assigned to a lineage, we set the ‘cutoff_method’ and ‘cor_threshold_static’ parameters of scClassify to ‘static’ and 0. In particular, to account for the limited number of data and the imbalance in classes in the training data, we performed bootstrapping of each of the reference datasets by randomly sampling 70% of the data to train the scClassify model. Furthermore, we addressed the class imbalance in the reference by randomly sampling with replacement to generate a balanced training data with class size matched to the median number of cells across the cell types. These strategies prevent the final model from becoming biased towards the majority classes and improving the model’s ability to accurately classify all cell lineages. The final cell types were determined as the most voted across 50 bootstraps, and the final classification probability was defined as the average of the 50 classification probabilities. Any cells that were not classified to any of the three lineages were labelled as unknown. Using the scClassify probabilities, we identified cells that have been assigned a cell type with high confidence (see “Proportion of high confidence cells“ of section “Derivation of fidelity metrics” for detailed description). Any cells labelled as an unknown or low confidence assignment were excluded from the finer developmental stage classification. Next, an additional scClassify model was then built to predict the developmental stage of the assigned cells. We built a model for each lineage using the finer level of cell type annotations that resolve the lineages into sub-states in terms of their developmental stage. As before, the cell type labels were determined by the majority voting from individual classification labels using each of the *in vivo* reference dataset.

#### Derivation of cell identity gene statistics from blastoids

To derive the cell identity gene statistics for the human blastoid datasets, the count matrix of cell-gene variables were normalised and log-transformed using the *logNormCounts* function from the scater package (McCarthy et al., 2017). The transformed and normalised data from each batch were subsequently analysed using the Cepo package (Kim et al., 2021) for quantifying cell identity gene statistics for each lineage resolved by time point based on the differential stability metric.

#### Benchmarking of the fidelity of human blastoids to the *in vivo* embryo reference

##### Derivation of fidelity metrics

To systematically compare the fidelity of protocols that generate blastoids to recapitulate the *in vivo* human embryo, we defined a panel of metrics that each assess different layers of transcriptomic similarity for comparing the blastoids. In total, we devised five major classes of evaluation metrics, of which four are resolved by lineages, leading to a total of 22 metrics.

**(1) Coverage:** We quantified the coverage of cell types generated from the blastoids through two metrics: cell lineage coverage and recapitulation of the cell heterogeneity in the *in vivo* reference. A blastoid was deemed to generate the cell lineage of interest if it contained >= 5 cells from that lineage with high confidence assignment. The coverage of the three lineages (epiblast, hypoblast, and trophoblast) was calculated as the sum of the three binary scores, and the coverage of the developmental stages was calculated as the sum of the binary scores for each lineage. Protocols with the capacity to generate all lineages (for the cell lineage coverage) and all the developmental stages (for the cell heterogeneity score) were assigned a scaled score of 1, whereas those with nil capacity to generate the expected cell types were assigned a score of 0. We considered a protocol that is able to generate more sub-states to better recapitulate the heterogeneity observed in the human embryo.
**(2) Proportion of high confidence lineage-specific cells:** To identify the high confidence cells, we used the final classification probabilities from scClassify. For each lineage, we extracted the probabilities for the cells originating for the lineage of interest. Then the cumulative sum was calculated on the log-transformed probabilities. We calculated cumulative sum of a sequence of log-transformed probabilities, *a*_1_, *a*_2_ … *a_n_*, where *a* is the log-transformed probability for a cell and *n* is the number of cells in a given lineage, using the following equation:

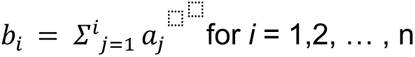 Where *b_i_* is the i-th element of the cumulative sum sequence and the summation symbol (∑) represents the sum of all numbers from *j*=1 to *i*. The cumulative sum of sequences were used to determine the point of inflection, which we defined as the first instance at which the difference between two elements in a sequence is greater than 0.01. Any cells that did not pass this threshold were considered as low confidence and were excluded from the benchmarking analysis. The final proportion of high confidence cells was defined as the following:

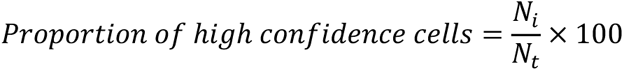

where *N_i_* represents the number of high confidence cells in lineage *i*, and *N_t_* represents the total number of cells in lineage *i*. Finally, the overall score was calculated by taking the arithmetic mean of the lineage-specific proportions.
**(3) Similarity of developmental stage-resolved cell identities against the embryo reference:** The lineage and developmental stage-specific fidelity of the blastoid was quantified using the pre- computed embryo reference. The reference consists of the top 500 Cepo statistics from each of three major lineages as well lineages that are further resolved by developmental stages. After obtaining the intersection of genes (which led to a total of 2,162 and 4,859 genes, respectively), we computed the developmental stage-resolved similarity of the blastoid data in terms of the Pearson’s correlation between the ranked Cepo statistics derived for the blastoid data and the corresponding ranked Cepo statistics from the reference. The overall cell identity score was calculated as the arithmetic mean of the lineage or developmental stage-resolved similarities. A higher score denotes stronger fidelity to the embryo reference and a lower score denotes weaker fidelity.
**(4) Lineage-to-lineage similarity:** The above two metrics are designed to quantify how well a given cell type found in the embryo reference is recapitulated in the blastoids. Whilst the metrics provide a quantitative measure for the predicted cell type found in the blastoid to its counterpart in the reference, it does not necessarily provide information about how well the relationships between a cell type with another cell type is captured. Therefore, we propose a metric called lineage-to-lineage similarity to dissect how well the between-lineage structure observed in the embryo reference is recapitulated by cells generated using the blastoid protocols, providing a measure of similarity on a cross-lineage scale that is distinct to the lineage-specific measures described above. Previously, we have shown that the cell identity scores computed from the Cepo statistics outperforms several existing marker gene detection tools to generate the known hierarchical lineage structures within large-scale atlas data (Kim et al., 2021). Thus, we used the Cepo statistics to compute the lineage-to-lineage structure by generating a pairwise matrix of lineage-to-lineage similarities where each row of the matrix depicts the relationship of the cell type of interest against all other cell types. We then compared these scores between the embryo reference and the blastoids in terms of the Pearson’s correlation. The overall score was calculated as the arithmetic mean of the lineage-to-lineage similarities. Again, a higher score denotes stronger fidelity to the embryo reference and a lower score denotes weaker fidelity.
**(5) Similarity to non-human primate embryos:** Because of the challenges in accessing human embryonic material that have not undergone extended *in vitro* culture, we evaluated the similarity in the cellular identities between the respective lineages from embryos derived from two non-human primates, the cynomolgus and the marmoset monkey, to further validate the blastoids. The embryos were harvested at Carnegie stages 3-6 for the cynomolgus monkey and at pre-implantation and Carnegie stages 5-7 for the marmoset monkey. We used a total of 175 cells (34 epiblast, 54 hypoblast, 87 trophoblast cells) from the cynomolgus embryo (Nakamura 2016, Nature) and 151 cells (38 epiblast, 53 hypoblast, 60 trophoblast cells) from the marmoset embryo (Boroviak 2018, Development). The processed data were downloaded using the accession numbers GSE74767 and E-MTAB-7078 for the cynomolgus and marmoset monkey data, respectively.

Limited to the resolution of the monkey data, we compared the broader lineage-level cellular identities (i.e., the epiblast, hypoblast, and the trophoblast) between the monkey embryos and the blastoids. To compare the cellular identities, we again followed the Cepo framework as in (3) and (4). In particular, we obtained the non-human primate cellular signature by taking the top 500 cell identity genes from each lineage and species. Using the intersection of the selected cell identity gene sets, we calculated Pearson’s correlation of the ranked Cepo statistics of these genes (165 genes in total) between the blastoid and the non-human primate embryos for each lineage. The overall score was calculated by taking the arithmetic mean of the lineage-specific scores. A higher correlation denotes a greater similarity to the non-human primate embryo, whilst a lower correlation denotes a weaker similarity.

##### Score aggregation and overall ranking of fidelity

To enable comparison of scores between different metric classes, we first normalised the individual metric scores through z-score transformation by shifting and scaling the scores to σ = 1 and μ = 0. We then performed min-max standardisation to bring the scores into the [0,1] range. To rank the protocols, we aggregated the scores by taking the rank of the protocols for each metric and then calculating the arithmetic mean across the different metrics. Finally, these benchmarking results were visualised using the funkyheatmap package that enables heatmap-like visualisations of benchmark data (Saelens et al., 2019).

